# Local autocrine signaling of IGF1 synthesized and released by CA1 pyramidal neurons regulates plasticity of dendritic spines

**DOI:** 10.1101/2021.04.08.439065

**Authors:** Xun Tu, Anant Jain, Helena Decker, Ryohei Yasuda

## Abstract

Insulin-like growth factor 1 (IGF1) regulates hippocampal plasticity, learning, and memory. While circulating, liver-derived IGF1 is known to play an essential role in hippocampal function and plasticity, IGF1 is also synthesized in multiple brain regions, including the hippocampus. However, little is known about the role of hippocampus-derived IGF1 in synaptic plasticity, the type of cells that may provide relevant IGF1, and the spatiotemporal dynamics of IGF1 signaling. Here, using a new FRET sensor for IGF1 signaling, we show that IGF1 in the hippocampus is primarily synthesized in CA1 pyramidal neurons and released in an activity-dependent manner in mice. The local IGF1 release from dendritic spines triggers local autocrine IGF1 receptor activation on the same spine, regulating structural and electrophysiological plasticity of the activated spine. Thus, our study demonstrates a novel mechanism underlying synaptic plasticity by the synthesis and autocrine signaling of IGF1 specific to CA1 pyramidal neurons.

## Main

Insulin-like growth factor 1 (IGF1) is a hormone that plays essential roles in cellular proliferation, differentiation, development, and metabolism in various tissues(Adams et al., 2000; Fernandez and Torres-Alemán, 2012; Ward and Lawrence, 2011). IGF1 binds and activates the IGF1 receptor (IGF1R), a receptor tyrosine kinase expressed widely in the brain(Kar et al., 1993), including excitatory neurons in the hippocampus(Gazit et al., 2016). The signaling pathway plays an essential role in the activity and plasticity of hippocampal synapses and hippocampus-dependent learning and memory(Cohen et al., 2009; Gazit et al., 2016; Stern et al., 2014; Trejo et al., 2007). Abnormal regulation of IGF1 in the brain is implicated in the cognitive decline in aging and brain diseases, including Alzheimer’s disease (AD)(Carro et al., 2002, 2006; Cohen and Dillin, 2008; Cohen et al., 2009; Moloney et al., 2010; Stern et al., 2014; Talbot et al., 2012; Trejo et al., 2007, 2001). It is known that circulating IGF1, which is primarily synthesized in the liver, enters the brain and contributes to hippocampal plasticity and cognitive functions(Nishijima et al., 2010; Trejo et al., 2007, 2001). IGF1 is also synthesized in multiple brain regions(Cao et al., 2011; Mardinly et al., 2016; Pristerà et al., 2019), including the hippocampus(Mitschelen et al., 2011a; Myhre et al., 2019; Sun et al., 2005a). However, it is unknown whether IGF1 synthesized in the hippocampus contributes to hippocampal synaptic plasticity, and if so, which types of cells synthesize relevant IGF1. It is also unknown whether IGF1 provides local synapse-specific signaling or global cell-wide signaling.

Here we show that hippocampal IGF1 is provided primarily by CA1 pyramidal neurons but much less in other types of cells in juvenile mice in the hippocampus. Furthermore, using a new FRET-based sensor for IGF1R activity, we found that IGF1 synthesized in CA1 pyramidal neurons is released from their dendritic spines in response to NMDA receptor activation to trigger local autocrine activation of IGF1R signaling on the same spines. The IGF1-IGF1R signaling in dendritic spines plays an essential role in regulating structural and electrophysiological plasticity. Thus, our study suggests the existence of the CA1-specific plasticity rule mediated by local IGF1 autocrine signaling.

### IGF1 receptor is required for structural plasticity of dendritic spines

To examine the role of IGF1-IGF1R signaling in the hippocampus, we deleted *Igf1r* in a sparse subset of neurons in organotypic hippocampal slices prepared from a conditional IGF1R knockout (*Igf1r*^fl/fl^) (Dietrich et al., 2000) by transfecting them with tdTomato-Cre and monomeric mEGFP (mEGFP) via a biolistic gene transfer (Harward et al., 2016). The *Igf1r* deletion altered neither the basal spine size nor the spine density in CA1 pyramidal neurons (**Supplementary Figure 1**). We induced structural plasticity in single dendritic spines on secondary or tertiary apical dendrites of CA1 pyramidal neurons by a train of two-photon glutamate uncaging pulses (0.5 Hz, 30 times) in the absence of extracellular Mg^2+^ (Harward et al., 2016; Matsuzaki et al., 2004). Under a control condition, in which tdTomato is expressed instead of tdTomato-Cre (Cre-), glutamate uncaging led to a spine volume increase lasting for more than ∼30 min (structural long-term potentiation or sLTP), as previously described (Harward et al., 2016; Matsuzaki et al., 2004) (**Figure 1a-c**). The volume increase occurred only in the stimulated spines, but not in surrounding spines. This structural plasticity is highly correlated with increases in synaptic strength and is considered the structural basis of electrophysiological long-term potentiation (LTP) (Matsuzaki et al., 2004). However, neurons transfected with tdTomato-Cre (Cre+) showed attenuated sLTP at a room temperature (24-26°C **; Figure 1a-c**) or a near-physiological temperature (31-32°C; **Supplementary Figure 2**). This deficit could be rescued by postsynaptic overexpression of mEGFP-tagged IGF1R (IGF1R-mEGFP). In addition, the bath application of specific inhibitors of IGF1R picropodophyllin (PPP; 5 µM) or PQ401 (20 µM) 30 min before sLTP induction inhibited sLTP (**Figure 1d,e**). These data demonstrate the critical role of postsynaptic IGF1R signaling in spine structural plasticity.

**Figure 1.**
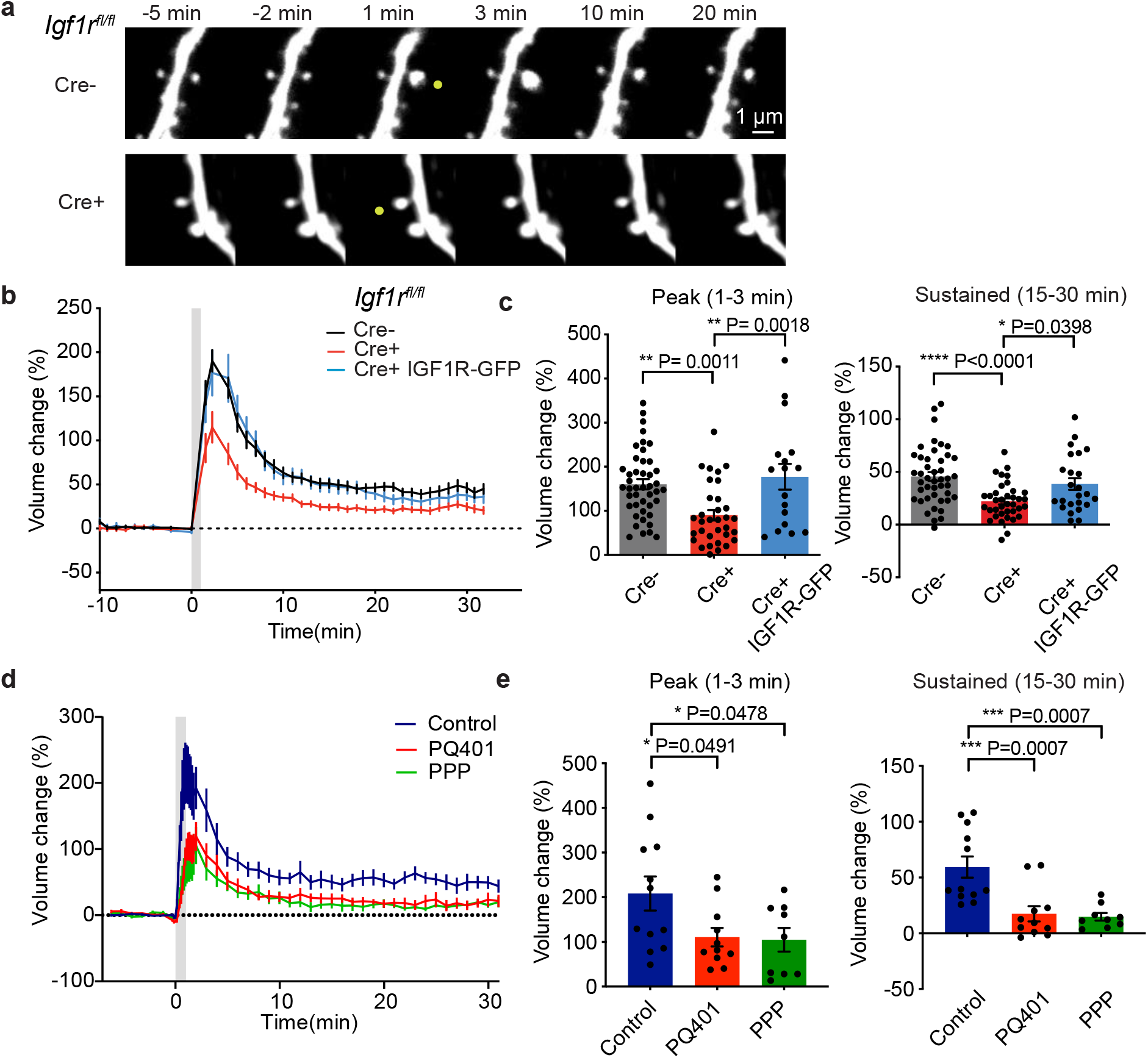
IGF1R is required for spine structural plasticity. **a-c**. Representative images (**a**), time course (**b**), and quantification (**c**) of glutamate-uncaging-induced spine volume change for *IGF1R*^fl/fl^ hippocampal slices in neurons transfected with (Cre+) or without Cre (Cre-) together with mEGFP, or IGF1R-mEGFP and Cre (Cre+ IGF1R-GFP). N = 44/17 for Cre+, 37/12 for Cre+, and 24/10 for Cre+ IGF1R-GFP (spines/cells). **d, e**. Time course (**d**) and quantification (**e**) of glutamate-uncaging-induced spine volume change for mouse hippocampal slices transfected with mEGFP in the presence of vehicle (Ctrl) or IGF1R inhibitors PQ401 or PPP. N = 12/5 for Ctrl, 11/5 for PQ401, and 9/5 for PPP (spines/cells). Data are means +/-s.e.m. Asterisks denote statistical significance (*p < 0.05, **p < 0.01, ***p < 0.001, ****p < 0.0001) as determined by ANOVA followed by a post-hoc test using Tukey’s correction.

### IGF1 receptor is locally activated in dendritic spines during structural plasticity

Given IGF1R is required for spine structural plasticity, we next asked whether the receptor in dendric spines is activated during this form of plasticity. To do so, we developed a fluorescence resonance energy transfer (FRET)-based sensor for IGF1R, optimized for two-photon fluorescence lifetime imaging (2pFLIM) (**Figure 2a**). Our IGF1R sensor consisted of two components: IGF1R-mEGFP and the SH2 domain of PI3K, an IGF1R binding partner, fused to two copies of mCherry (mCherry-SH2-mCherry). When IGF1R is auto-phosphorylated at Y1316 upon its activation, mCherry-SH2-mCherry binds to IGF1R-mEGFP, increasing FRET between EGFP and mCherry. Overexpressed IGF1R-mEGFP function similarly to endogenous IGF1R, as it rescued the impaired sLTP caused by IGF1R knockout (**Figure 1b, c**). The sensor expressed in HeLa cells produced FRET signals sensitive to IGF1, specific for Y1316 phosphorylation, and reversible by IGF1R inhibitor (**Supplementary Figure 3**). The sensor expressed in CA1 pyramidal neurons in organotypic hippocampal slices of mice showed an increase in FRET by bath application of IGF1 in a dose-dependent manner, with a dissociation constant similar to reported values (*K*_d_ = 0.61 nM; reported values: 0.16-0.43 nM) (**Figure 2b, c**) (Adams et al., 2000; Forbes et al., 2002).

**Figure 2.**
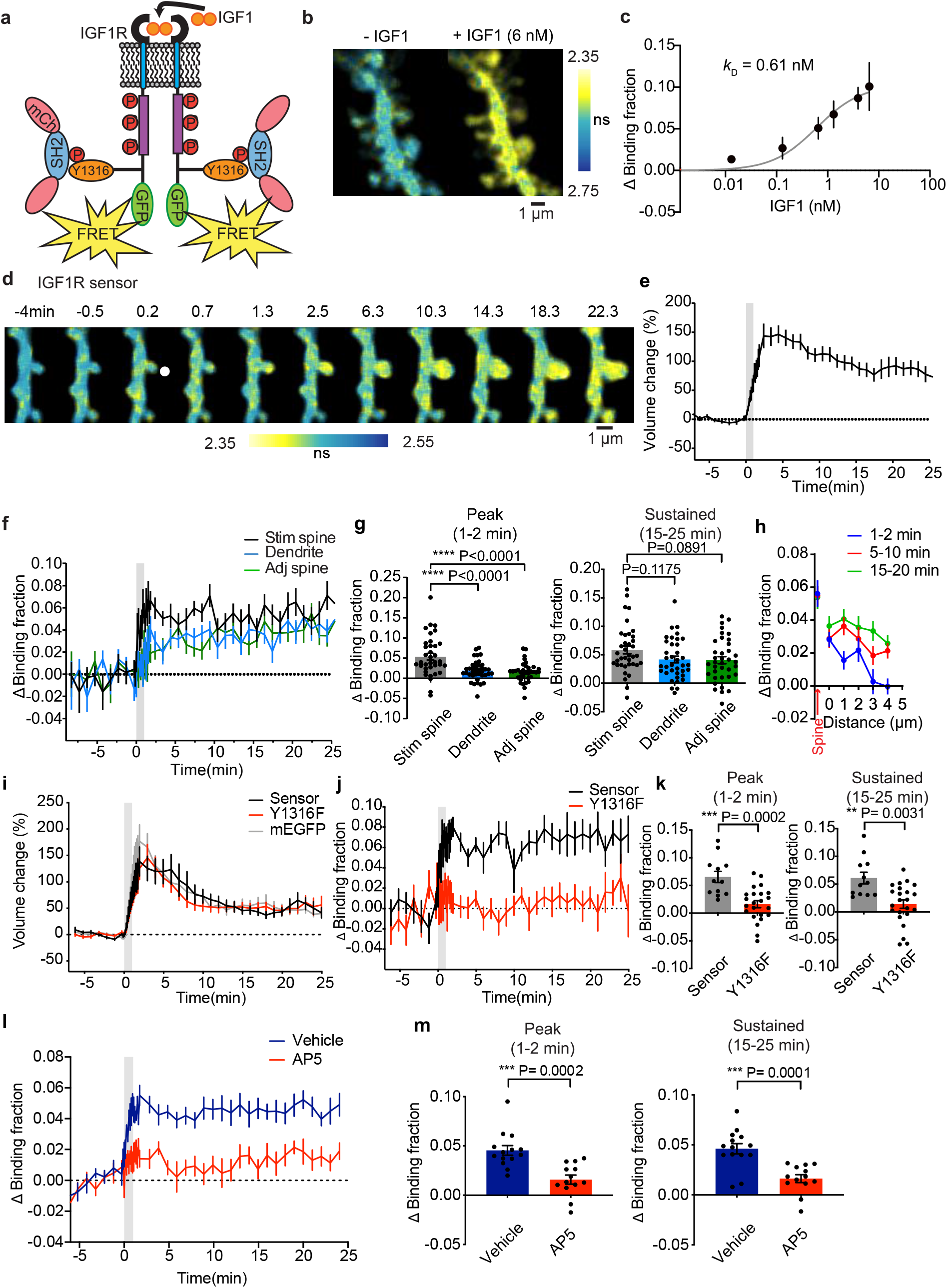
Glutamate uncaging induces rapid, local, and persistent IGF1R activation. **a**. IGF1R sensor design. GFP: mEGFP, RFP: mCherry, SH2: SH2 domain of p85a, an IGF1R effector. **b, c**. Representative images (**b**) and quantification (**c**) of IGF1R activation, measured as the change in sensor binding fraction in response to a different dose bath IGF1. The Michaelis-Menten curve (*K*_d_ = 0.61 µM; obtained by fitting) is also shown in **c**. N = 20 (neurons). **d**. Representative 2pFLIM images of IGF1R activation averaged over indicated time points. Arrowhead represents the uncaging location. Shorter lifetime means higher IGF1R activity. **e-g**. Volume change for the stimulated spine (**e**), and time course (**f**), and quantification (**g**) of IGF1R activation in stimulated spines (Stim spine), adjacent spines (Adj spine), and their parent dendritic shaft (Dendrite). N = 37/18 for stimulated spines and dendritic shafts, and 35/18 for adjacent spines (spines/cells). **h**. Spatial profile of IGF1R activation, measured as the change in the binding fraction of the dendrite as a function of distance from the stimulated spine. N = 36/18 (stimulated spines/cells) **i-k**. Spine volume change (**i**), time course (**h**), and quantification (**k**) of IGF1R activation in neurons expressing the IGF1R sensor (sensor; separate data from **d-g**) or a <u>mutant</u> sensor in which the critical phosphorylation site is mutated (Y1316F). For volume change, data from neurons expressing mEGFP (**Figure 1d**) are also presented for comparison in **i**. N = 12/6 for sensor and 23/12 for Y1316F (spines/cells). **l, m**. Time course (**l**) and quantification (**m**) of IGF1R activation in the presence of vehicle (water), or AP5 (100 µM). Spine volume changes for this data are in **Supplementary Figure 6**. N = 13/7 for vehicle, and N= 13/7 for AP5 (spines/cells). Data are means +/- s.e.m. Asterisks denote statistical significance (*p < 0.05, **p < 0.01, ***p < 0.001, ****p < 0.0001) as determined by a two-tailed t-test (**k, m**) or ANOVA followed by a post-hoc test using Tukey’s correction (**g**).

Next, we investigated IGF1R activity during spine structural plasticity in CA1 pyramidal neurons expressing our IGF1R sensor (**Figure 2d**). In response to glutamate uncaging targeted to a single spine, IGF1R was rapidly activated in the stimulated spine, peaking at ∼1–2 min and remaining elevated for at least 25 min (**Figure 2d-f**). The peak amplitude was ∼60% saturation level (0.59 ± 0.06), corresponding to ∼0.9 nM. The IGF1R activation was restricted to the stimulated spine during the first 30–60 s after the onset of glutamate uncaging but slowly spreads over a few minutes over a few micrometers along the parent dendrite, invading adjacent spines (**Figure 2g, h**). A mutant sensor, in which we mutated the IGF1R phosphorylation site critical for interacting with the SH2 domain (Y1316F), abolished the FRET signal change in response to glutamate uncaging, suggesting the specificity of the measurement (**Figure 2i-k**). Spine volume change in neurons expressing the sensor or Y1316F sensor was indistinguishable from neurons expressing mEGFP, implying that the sensor expression has a minimum impact on spine plasticity (**Figure 2i**). The spatiotemporal profile of IGF1R activation was at a near-physiological temperature similar to that at a room temperature (**Supplementary Figure 4**). The amplitude of the sensor response and spine volume change did not correlate with the sensor’s expression level (**Supplementary Figure 5)**. Inhibiting the NMDA receptor with AP5 (100 µM) abolished IGF1R activation (**Figure 2l, m**) as well as sLTP (**Supplementary Figure 6**), indicating that IGF1R activation requires the activation of NMDA receptors. Finally, IGF1R activations in females and males were similar (**Supplementary Figure 7**).

### Synthesis, storage, and secretion of IGF1 in CA1 hippocampal neurons

To address the source of IGF1 that activates IGF1R in the dendritic spines, we examined IGF1 expression in the hippocampus by immunostaining brain sections of juvenile mice (P20) with an anti-IGF1 antibody (**Figure 3a-d** for conditional knockout of IGF1 (*Igf1*^fl/fl^), **Supplementary Figure 8** for wildtype mice). We found a punctuated pattern primarily in dendrites and soma specifically for CA1 pyramidal neurons, but little for other types of cells, including CA3 pyramidal neurons (**Figure 3a; Supplementary Figure 8**). To verify the specificity of the antibody signal and IGF1 synthesis in CA1 pyramidal neurons, we removed *Igf1* from a sparse subset of excitatory neurons in *Igf1*^fl/fl^ mice by in-utero injection of AAV-Camk2a-Cre and AAV-CAG-Flex-EGFP (Harward et al., 2016). We found that the IGF1 signals in Cre-expressing, EGFP-expressing cells (Cre+) were significantly lower than that in nearby cells without EGFP expression (Cre-) for CA1 pyramidal neurons (**Figure 3b, d**). In contrast, CA3 pyramidal neurons showed much lower IGF1 signals both in Cre+ and Cre-cells (**Figure 3b, d**). EGFP expression with AAV-CAG-EGFP without Cre did not change IGF1 in CA1 pyramidal neurons (**Figure 3c, d**). These results indicate that the immunofluorescence signal in CA1 pyramidal neurons is mostly from IGF1 synthesized in the same neurons, and other cell types in the hippocampus express much less IGF1.

**Figure 3.**
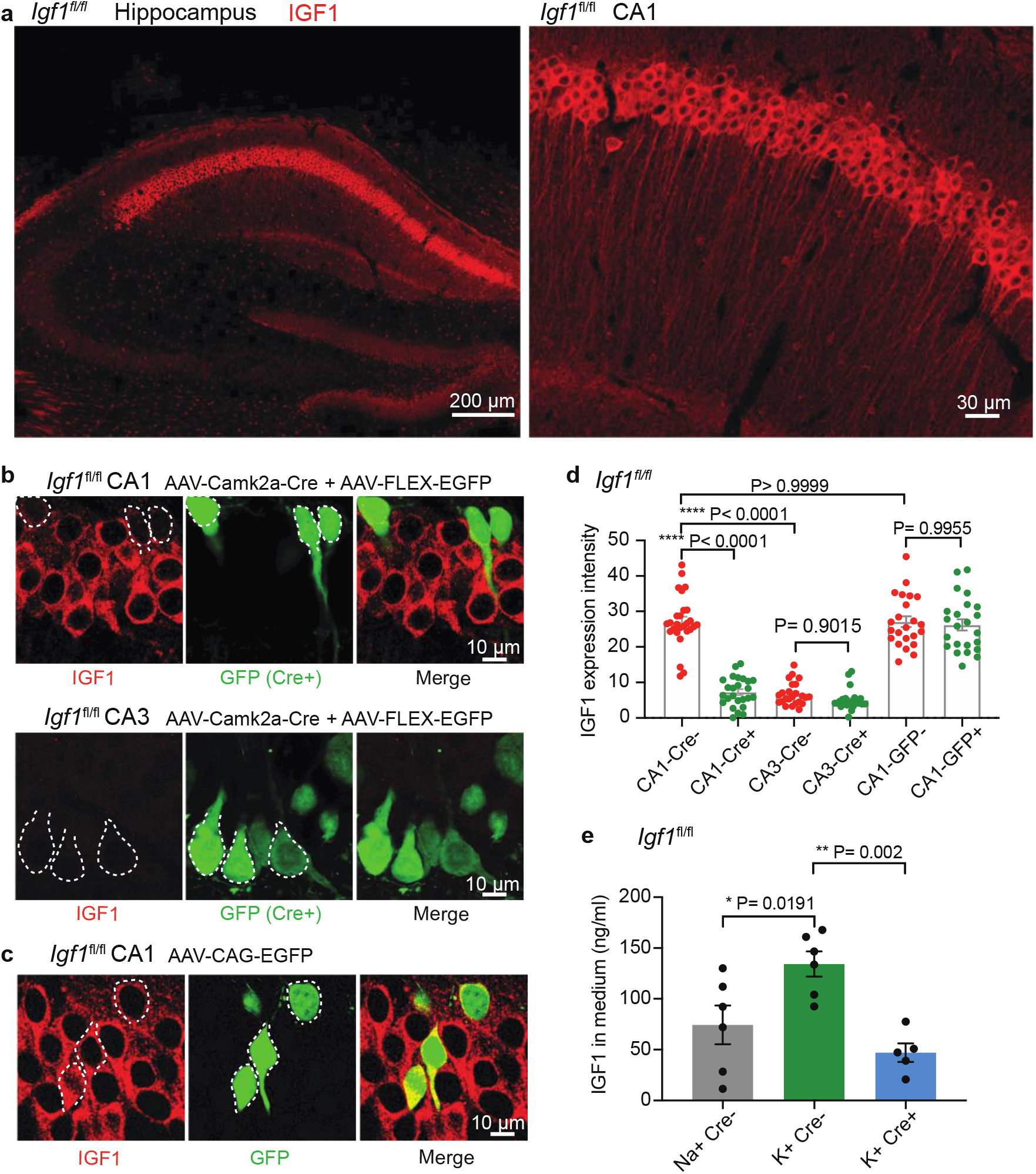
Synthesis and activity-dependent secretion of IGF1 by CA1 pyramidal neurons. **a**. Immunofluorescence images of the entire hippocampus and CA1 region of *Igf1*^fl/fl^ mice (P20) stained with an anti-IGF1 antibody. **b**. IGF1 immunofluorescence images of CA1 and CA3 neurons in *Igf1*^*fl/fl*^ mice transduced with AAV-Camk2a-Cre together with AAV-CAG-EGFP by in-utero intraventricular injection. *Igf1* is deleted in Cre+, EGFP expressing neurons. **c**. IGF1 immunofluorescence images of CA1 neurons in *Igf1*^*fl/fl*^ mice transduced with AAV-CAG-EGFP without Cre. **d**. Quantification of immunofluorescence intensity of IGF1 in Cre+ and surrounding Cre-pyramidal neurons for CA1 or CA3 (condition in **b**), and in EGFP+ and surrounding EGFP-(without Cre) in CA1 pyramidal neurons (condition in **c**). **e**. Activity-dependent secretion of IGF1 from organotypic slice culture prepared from *Igf1*^fl/fl^ mice. IGF1 secretion was quantified by ELISA assay on culture medium after 24 h incubation in serum-free medium plus 20 mM NaCl or 20 mM KCl. The slices were transduced either with AAV-CMV-LacZ (NaCl Cre-, KCl Cre-) or AAV-Camk2a-Cre (KCl Cre+). Data are means +/- s.e.m. Asterisks denote statistical significance (*p < 0.05, **p < 0.01, ****p < 0.0001) as determined by ANOVA followed by a post-hoc test using Tukey’s correction.

Given the CA1 pyramidal neuron is a primary source of IGF1 in the hippocampus, we asked whether these neurons can release synthesized IGF1 by measuring extracellular IGF1 in the medium from organotypic hippocampal slices prepared from *Igf1*^fl/fl^ mice with ELISA assay (**Figure 3e**). We found that the medium from control slices, transduced with AAV-CMV-LacZ, incubated in high potassium solution (+20 mM) over 24 h contained significantly higher IGF1 than that in high sodium solution (+20 mM), suggesting that IGF1 release is activity-dependent. Furthermore, *Igf1* deletion from excitatory neurons with AAV-Camk2a-Cre significantly reduced IGF1 in the medium, indicating that IGF1 in the medium was primarily synthesized and released by excitatory neurons.

### Postsynaptic synthesis of IGF1 is necessary for IGF1R activation and structural and electrophysiological spine plasticity

Next, we examined whether IGF1 synthesized in CA1 pyramidal neurons contributes to sLTP using hippocampal slices prepared from *Igf1*^fl/fl^ transfected sparsely with tdTomato-Cre and mEGFP (Liu et al., 1998). Deleting *Igf1* did not cause detectable effects on basal spine density and morphology, compared with control neurons expressing tdTomato instead of tdTomato-Cre (**Supplementary Figure 9**). However, it attenuated glutamate-uncaging-evoked sLTP at room temperature (**Figure 4a, b**) or a near-physiological temperature (**Supplementary Figure 10**), suggesting that postsynaptically synthesized IGF1 is required for sLTP. Bath application of a saturating concentration of IGF1 (6.5 nM) rescued sLTP for the sustained phase but not for the transient phase. We also measured IGF1R activity in IGF1 knockout cells by biolistically overexpressing the IGF1R FRET sensor together with tdTomato-Cre in *Igf1*^fl/fl^ mice (**Figure 4c**). The *Igf1* deletion significantly inhibited IGF1R activation in the stimulated spines in these neurons, suggesting that postsynaptically synthesized IGF1 contributes to the IGF1R activation in spines on the same neuron, likely through local IGF1 release from the same spine (**Figure 4c, d; Supplementary Figure 11**).

**Figure 4.**
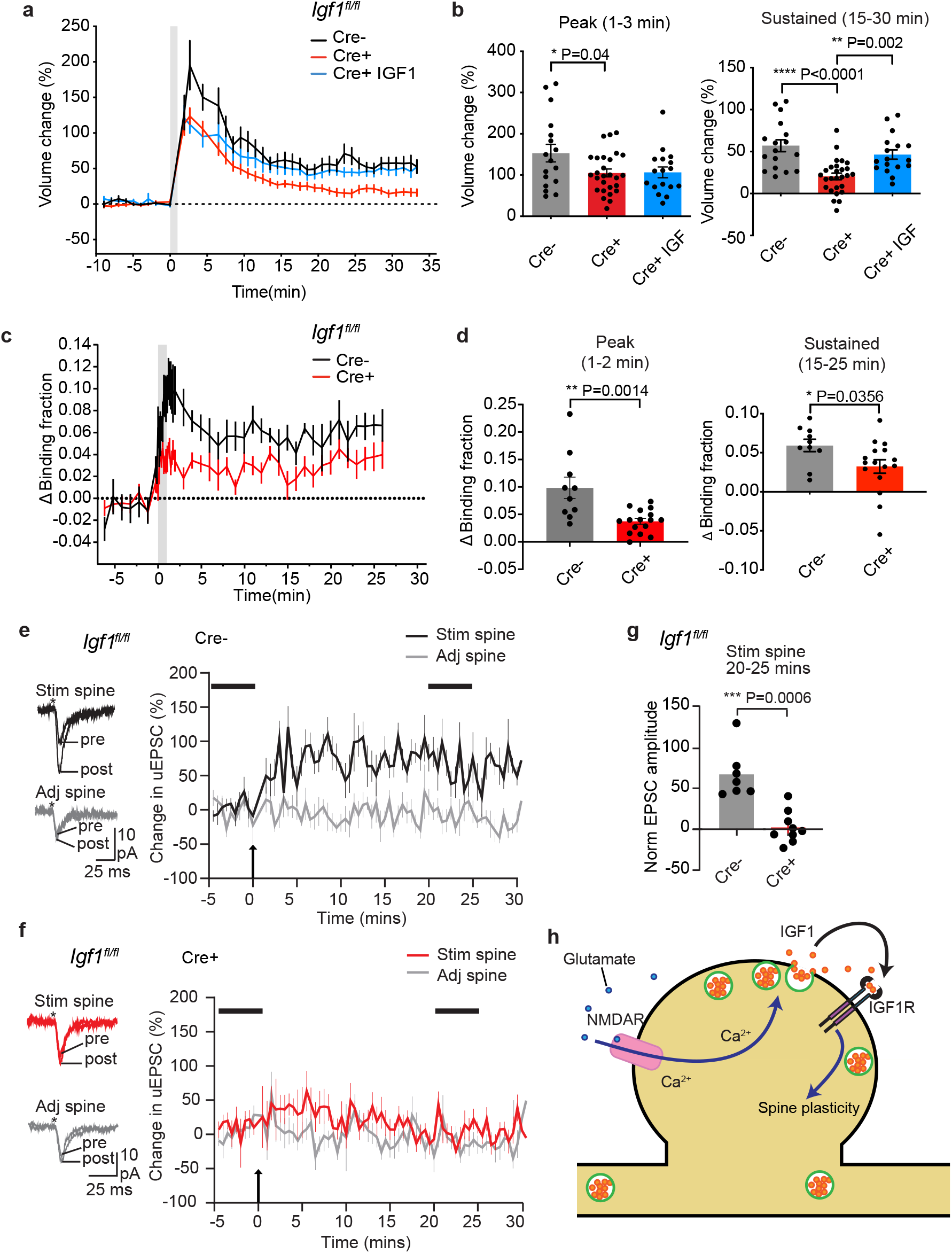
Postsynaptically synthesized IGF1 is required for spine plasticity and IGF1R activation. **a, b**. Time course (**a**) and quantification (**b**) of glutamate-uncaging-induced spine volume change for *Igf1*^fl/fl^ hippocampal slices in neurons transfected with (Cre+) or without Cre (Cre-). For Cre+ IGF1, Cre positive cells were treated with IGF1 (6.5 nM) for 10 min before glutamate uncaging. N = 17/7 for Cre-, 27/13 for Cre+, and 17/8 for Cre+ IGF1 (spines/cells). **c, d**. Time course (**c**) and quantification (**d**) of glutamate-uncaging-induced IGF1R activation for *Igf1*^fl/fl^ hippocampal slices in neurons transfected with (Cre+) or without Cre (Cre-). N = 10/6 for Cre- and 16/8 for Cre+. Spine volume changes for these data are in **Supplementary Figure 11.** **e, f**. Raw trace of uncaging-evoked postsynaptic currents (uEPSCs) averaged over 10 trials before (−5 to 0 min) and after (20 to 25 min) LTP induction by pairing depolarization (0 mV) with single spine uncaging (1 Hz, 20 times) (right), and the time courses of changes in the peak uEPSC amplitude (left) in CA1 pyramidal neurons in *Igf1*^fl/fl^ organotypic slices transduced with AAV-Camk2a-Cre and AAV-CAG-EGFP. Recordings from Cre-(EGFP-negative) (**e**) or Cre+ (EGFP positive) neurons (**f**). **g**. Quantification of LTP for Cre+ and Cre- neurons (**e, f**) as measured as the ratio of uEPSCs averaged over 10 trials before (−5 to 0 min) and after (20 to 25 min) LTP induction. N = 9/7 for Cre+, 7/6 for Cre-(Neurons/Slices). **h**. Schematic model of IGF autocrine signaling in the dendritic spine. Data are means +/-s.e.m. Asterisks denote statistical significance (*p < 0.05, **p < 0.01, ***p < 0.001, ****p < 0.0001) as determined by a two-tailed t-test (**d, g**) or an ANOVA followed by a post-hoc test using Tukey’s correction (**b**).

To further assess whether postsynaptically synthesized IGF1 is required for electrophysiological LTP, we performed a whole-cell patch-clamp on neurons in *Igf1*^fl/fl^ slices transduced with AAV-Camk2a-Cre together with AAV-Flex-EGFP (1:15 – 1:100). Under this condition, only Cre positive neurons, in which *Igf1* is deleted, are highlighted with mEGFP (Cre+). We paired postsynaptic depolarization (0 mV) with 2-photon glutamate uncaging (1 Hz, 20 times) to induce LTP at a single spine (Matsuzaki et al., 2004). In control neurons (Cre-), normal LTP was induced, as indicated by a potentiation in uncaging-evoked postsynaptic currents (uEPSCs) in stimulated spines but not in surrounding spines (**Figure 4e-g**). However, in Cre+ neurons, we failed to induce LTP (**Figure 4e-g**), suggesting that IGF1 synthesized in the postsynaptic neuron is required for LTP.

## Discussion

Overall, this study demonstrates that IGF1 is synthesized and secreted by CA1 pyramidal neurons and plays critical roles in synaptic plasticity through local autocrine signaling in single dendritic spines (**Figure 4h**). Immunostaining data, combined with *Igf1* deletion in a small subset of pyramidal neurons, suggest that most IGF1 in the hippocampus is synthesized and stored in CA1 pyramidal neurons, but much less in other types of cells in mice. Our imaging data indicate that the synthesized IGF1 is released from dendritic spines in an activity-dependent manner and triggers autocrine IGF1R activation in the same dendritic spines. This local autocrine IGF1-IGF1R signaling is essential for structural and electrophysiological plasticity of dendritic spines. Our IGF1R activation measurements suggest that transient IGF1 elevation at the synapse during spine structural plasticity reaches ∼0.9 nM. Although the local resting concentration of free IGF1 *in vivo* is yet to be determined, IGF1 concentration in the hippocampus homogenate has been reported to be relatively low (0.16-0.8 nM)(Adams et al., 2009; Gomes et al., 2009; Mitschelen et al., 2011b). Since a saturating concentration of bath IGF1 (6.5 nM) only partially rescues the plasticity phenotype of postsynaptic *Igf1* deletion, spine plasticity may specifically require the local IGF1 released at the synapse. Bulk IGF1 supplied from the circulation or released by surrounding cells may play an additional role in synaptic plasticity (Myhre et al., 2019; Sun et al., 2005b; Trejo et al., 2001). Overall, our study indicates that one primary source of IGF1 relevant to hippocampal plasticity is those synthesized and released by the CA1 pyramidal neuron. Since the presynaptic function requires IGF1R activity(Gazit et al., 2016), the postsynaptically released IGF1 may simultaneously regulate post-and presynaptic functions. Finally, impaired regulation of IGF1 synthesis and release by CA1 pyramidal neurons may cause an abnormal IGF1 level in the hippocampus, potentially leading to the impairment of hippocampal plasticity in aging and brain diseases(Carro et al., 2002, 2006; Cohen and Dillin, 2008; Cohen et al., 2009; Moloney et al., 2010; Talbot et al., 2012; Trejo et al., 2007, 2001).

## Acknowledgments

We would like to thank Tavita Garret for pilot experiments in the initial phase of the project. Drs. Lesley Colgan, Paula Parra-Bueno and Sarah Stern for critical reading; Dr. Inna Slutsky for the IGF1R plasmid; Minida Dowdy, Elizabeth Garcia, and the MPFI ARC for animal care; members of the Yasuda laboratory for discussion; Dr. Long Yan for helping with the microscope; and David Kloetzer for managing the laboratory. This work was funded by Louis D Srybnik Foundation Inc. & Foundation for the Art, Science, and Education Inc, NIH (R35NS116804, DP1NS096787, R01MH080047), and the Max Planck Florida Institute for Neuroscience.

## Author contributions

XT, HD, RY conceptualized the project, XT acquired and analyzed most of the data. AJ acquired and analyzed electrophysiology data, XT, RT wrote the manuscript. All authors discussed and agreed on the content of the manuscript.

## Materials and Methods

### Reagents

Human recombinant IGF1 was purchased from Thermo Fisher, and K252a, d-2-amino-5-phosphonovalerate (d-AP5), PQ401, and Picropodophyllin (PPP) were from Tocris.

### Animals

All experimental procedures were approved and carried out in accordance with the regulations of the Max Planck Florida Institute for Neuroscience Animal Care and Use Committee in accordance with guidelines by the US National Institutes of Health. P4-P8 mouse pups from both sexes were used for organotypic slices for imaging studies. C57BL/6, *Igf1r*^fl/fl^ (#012251), and *Igf1*^fl/fl^ (#016831) mice were from The Jackson Laboratory.

### Plasmids

Our IGF1R sensor consists of two components, IGF1R-mEGFP and mCherry-SH2-mCherry. IGF1R-mEGFP was prepared by inserting the coding sequence of IGF1R (a gift from Dr. Inna Slutsky) into pEGFP-N1 (Clontech) containing the A206K monomeric mutation in EGFP and the CAG promoter. The linker between IGF1R and mEGFP is GGSGGS. mCherry-SH2-mCherry was prepared from the C-terminal SH2 domain of p85a (Addgene #46463). The linker between mCherry and SH2 is SGLRSRAQASNSAVDGTA for the N terminus and GSG for the C terminus.

### HeLa cell maintenance, transfection and imaging

HeLa cells (ATCC CCL-2) were cultured in Dulbecco’s modified Eagle medium (DMEM) supplemented with 10% FBS at 37 °C in 5% CO2. Cells were transfected with Lipofectamine 2000 using the manufacturer’s protocol (Invitrogen). For sensor validation, the IGF1R sensor was transfected with the plasmid ratio of 1:4 (donor:acceptor). Then, 24–48 h later, culture media was replaced with insulin- and serum-free DMEM media. Imaging was performed 2 h after exclusion of insulin and serum in HEPES-buffered ACSF (HACSF; 20 mM HEPES, 130 mM NaCl, 2 mM NaHCO3, 25 mM d-glucose, 2.5 mM KCl and 1.25 mM NaH2PO4; adjusted to pH 7.4 and 310 mOsm) using 2pFLIM as described below. Cells were stimulated by adding IGF1 (6.5 nM) or vehicle to the bath.

### Organotypic hippocampal slice cultures and transfection

Organotypic hippocampal slices were prepared from wildtype or transgenic postnatal 4-8 day old mouse pups of both sexes as previously described (Stoppini et al., 1991). In brief, the animals were anesthetized with isoflurane, after which the animal was quickly decapitated and the brain removed. The hippocampi were dissected and cut into 350 µm thick coronal hippocampal slices using a McIlwain tissue chopper (Ted Pella, Inc) and plated on hydrophilic PTFE membranes (Millicell, Millipore) fed by culture medium containing MEM medium (Life Technologies), 20% horse serum, 1mM L-Glutamine, 1mM CaCl_2_, 2mM MgSO_4_, 12.9mM D-Glucose, 5.2mM NaHCO_3_, 30mM Hepes, 0.075% Ascorbic Acid, 1 µg/ml insulin. The slices were incubated at 37 °C in 5% CO^2^. After 7-12 days in culture, CA1 pyramidal neurons were transfected with biolistic gene transfer (O’Brien and Lummis, 2006) using 1.0 µm gold beads (8–12 mg) coated with plasmids containing cDNA of interest in the following ratios. GFP+tdTomato: 25ug GFP, 25ug tdTomato; GFP+Cre-tdTomato: 25ug GFP, 25ug Cre-tdTomato; IGF1R sensor: 12.5ug IGF1R-mEGFP, 37.5ug mCherry-SH2-mCherry; IGF1R sensor with tdTomato: 10ug IGF1R-mEGFP, 30ug mCherry-SH2-mCherry, 10ug tdTomato; IGF1R sensor with Cre-tdTomato: 10ug IGF1R-mEGFP, 30ug mCherry-SH2-mCherry, 10ug Cre-tdTomato. Neurons expressing mEGFP were imaged 2–7 days after transfection. Neurons expressing the IGF1R sensor were imaged 2-5 days after transfection. For conditional knockout experiments, imaging was performed 7-9 days after transfection to ensure the deletion of the target gene.

### 2pFLIM

FLIM imaging using a custom-built two-photon fluorescence lifetime imaging microscope was performed as previously described (Murakoshi et al., 2011). 2pFLIM imaging was performed using a Ti-sapphire laser (Coherent, Cameleon) at a wavelength of 920 nm with a power of 1.4-1.6 mW. Fluorescence emission was collected using an immersion objective (60×, numerical aperture 0.9, Olympus), divided with a dichroic mirror (565 nm), and detected with two separated photoelectron multiplier tubes placed after wavelength filters (Chroma, 510/70-2p for green and 620/90-2p for red). Both red and green channels were fit with photoelectron multiplier tubes (PMT) having a low transfer time spread (H7422-40p; Hamamatsu) to allow for fluorescence lifetime imaging. Photon counting for fluorescence lifetime imaging was performed using a time-correlated single-photon counting board (Time-harp 260, Pico-Quant) using custom software developed in C# (Pologruto et al., 2003) (https://github.com/ryoheiyasuda/FLIMage_public). 2pFLIM images were collected at 128×128 pixels at the frame rate of 7.8-Hz and averaged over 24 frames. Relative spine volume was measured as integrated fluorescence intensity of the spine (we used mEGFP fluorescence for neurons expressing mEGFP and Cre, and mCherry fluorescence for neurons expressing IGF1R sensor).

### Two-photon glutamate uncaging

A second Ti-sapphire laser tuned at a wavelength of 720 nm was used to uncage 4-methoxy-7-nitroindolinyl-caged-l-glutamate (MNI-caged glutamate) with a train of 4–8 ms, 2.8-3.0 mW pulses (30 times at 0.5 Hz) located ∼0.5 µm away from the spine of interest as previously described (Colgan et al., 2018). Experiments were performed in Mg^2+^ fee artificial cerebral spinal fluid (ACSF; 127 mM NaCl, 2.5 mM KCl, 4 mM CaCl_2_, 25 mM NaHCO_3_, 1.25 mM NaH_2_PO_4_ and 25 mM glucose) containing 1 µM tetrodotoxin (TTX) and 4 mM MNI-caged L-glutamate aerated with 95% O_2_ and 5% CO_2._ Experiments were performed at room temperature (24-26 °C) or 30-32 °C.

#### 2pFLIM analysis

To measure the fraction of the donor that was undergoing FRET with the acceptor (binding fraction), we fit a fluorescence lifetime curve summing all pixels over a whole image with a double exponential function convolved with the Gaussian pulse response function:

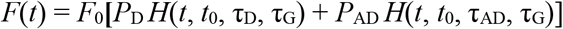

where τ_AD_ is the fluorescence lifetime of the donor bound with the acceptor, *P*_D_ and *P*_AD_ are the fraction of free donor and donor undergoing FRET with the acceptor, respectively, and *H*(*t*) is a fluorescence lifetime curve with a single exponential function convolved with the Gaussian pulse response function:

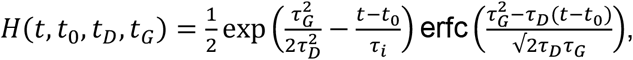

in which τ_D_ is the fluorescence lifetime of the free donor, τ_G_ is the width of the Gaussian pulse response function, *F*_0_ is the peak fluorescence before convolution and *t*_0_ is the time offset, and erfc is the complementary error function.

We fixed τ_D_ to the fluorescence lifetime obtained from free mEGFP (2.6 ns) and then fixed τ_AD_ to fluorescence lifetime of the donor bound with the acceptor (1.1 ns). For experimental data, we fixed τ_D_ and τ_AD_ to these values to obtain stable fitting.

To generate the fluorescence lifetime image, we calculated the mean photon arrival time, <*t*>, in each pixel as:

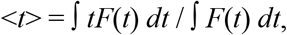

Then, the mean photon arrival time is related to the mean fluorescence lifetime, <*τ*>, by an offset arrival time, *t*_*o*_, which is obtained by fitting the whole image:

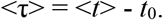

For small regions-of-interest (ROIs) in an image (spines or dendrites), we calculated the binding fraction (P_AD_) as:

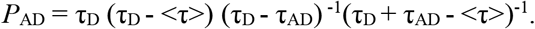

Data with baseline lifetime fluctuations greater than 0.15 ns were excluded before further analysis. To analyze the spatial spreading of IGF1 activation, contiguous 1 μm diameter ROIs along dendrite from the base of the stimulated spines were analyzed for IGF1 activation over time.

### In utero viral injection for single-cell IGF1 knockout

E14.5/15.5 timed-pregnant *Igf1*^fl/fl^ mice were deeply anesthetized using an isoflurane–oxygen mixture. The uterine horns were exposed, and approximately 1–2 μl of AAV solution mix (containing a mixture of AAV-CAG-Flex-EGFP at 6 × 10^12^ viral genome (vg)/ml and AAV-Camk2a-Cre at 2 × 10^12^ vg/ml or AAV-CAG-EGFP at 2 × 10^12^ vg/ml, all from Addgene) was injected through a pulled-glass capillary tube into the right lateral ventricle of each embryo.

### Measurements of IGF-1 Secretion

Organotypic hippocampal slices prepared from *Igf1*^*fl/fl*^ mice were transduced with AAV-CMV-LacZ or AAV-Camk2a-Cre after 4-5 days in culture. 8-10 days after virus infection, the slices were incubated for 1 day in a fresh tissue medium (without insulin/serum) containing either 20 mM NaCl or KCl, with adjusted osmolarity. After stimulation, the IGF-1 concentration in the medium was measured using the Mouse/Rat IGF-I/IGF-1 Quantikine ELISA Kit (R&D Systems, Inc., Minneapolis, MN, USA)

### Electrophysiology

Organotypic hippocampal slices prepared from *Igf1*^*fl/fl*^ mice were transduced with AAV-Camk2a-Cre and AAV-Flex-EGFP (1:15 – 1:100). Whole-cell patch-clamp was performed on Cre+ (EGFP positive) or Cre-cells (EGFP negative) in ACSF containing 4 mM MNI caged glutamate, 1 mM MgCl_2_ and 2 mM CaCl_2_ and 1 µM TTX with a patch pipette including an internal solution (145 mM K gluconate, 14 mM phosphocreatine, 4 mM NaCl, 0.3 mM NaGTP, 4 mM MgATP, 3 mM L-ascorbic acid, 50 µM Alexa-594, and 10 mM HEPES, pH 7.4, 315 mOsm). The current was measured under the voltage-clamp mode by a patch-clamp amplifier (MC-700B, Molecular Devices) and digitizer (National Instruments). The membrane potential was held at –70 mV. After a few minutes of dye loading, fluorescence from Alexa-594 was used to find dendritic spines in 2pFLIM. Uncaging-evoked postsynaptic currents (uEPSCs) were induced by photolyze caged glutamate at the interval of 30 s with 720 nm laser located near a spine, ∼0.5µm away from the tip of the spine. The uEPSC amplitude was 8-21 pA. LTP was induced by pairing depolarization of the cell to 0 mV and uncaging at 1 Hz, 20 times. Experiments were performed at room temperature (24-26 °C).

### Statistical analysis

All values are presented as mean ± SEM unless otherwise noted. The number of independent measurements (n[neurons/spines]) is indicated in Figure legends. Unpaired two-tailed student’s t-test was used for comparing two independent samples. Two-way ANOVA, followed by multiple comparison tests, was used to compare grouped data sets (Prism 6, GraphPad). Data were excluded if obvious signs of poor cellular health (for example, dendritic blebbing, spine collapse) were apparent.

---

**Supplementary Figure 1.**
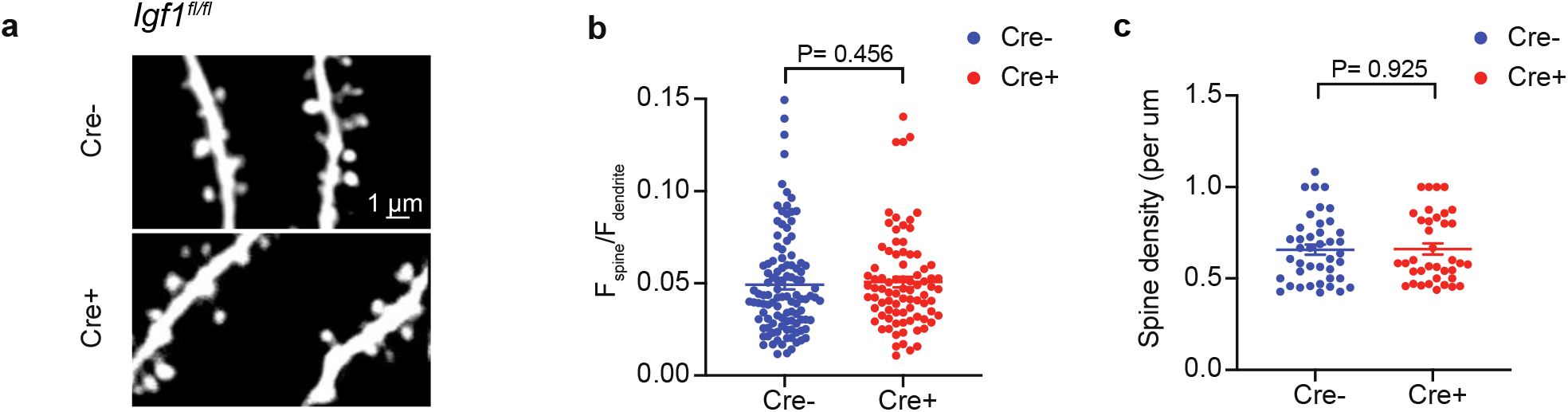
Deletion of IGF1R does not alter basal spine size and density. **a**. Representative 2-photon images of spines in neurons transfected with mEGFP plus tdTomato (Cre-) or mEGFP plus toTomato-Cre (Cre+) for *Igf1r*^*fl/fl*^ slices. **b**. The ratio of fluorescence intensity in the spine and that in the dendrite (sum intensity of spine divided by sum intensity of dendrite per 5 µm). N=108/8 for Cre- and N=83/7 Cre+ (spines/cells). **c**. Density of spines. N=41/8 for Cre- and N= 37/7 Cre+ (dendrites/cells). Data are means +/- s.e.m. P values are determined by a Mann-Whitney U test (**B**) or a two-tailed t-test (**C**).

**Supplementary Figure 2.**
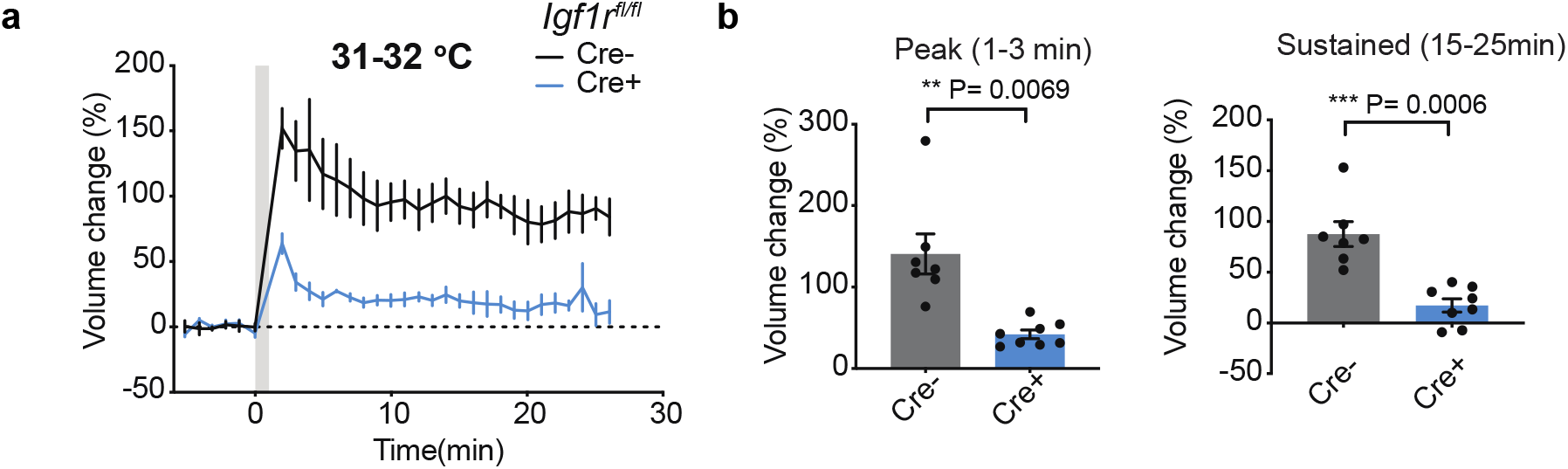
Spine structural plasticity is IGF1R-dependent at a near-physiological temperature. Time course (**a**) and quantification (**b**) of glutamate-uncaging-induced spine volume change for *Igf1r*^fl/fl^ hippocampal slices in neurons transfected with (Cre+) or without Cre (Cre-) together with mEGFP at 31-32 °C. N= 7/4 for Cre- and N= 8/4 for Cre+ (spines/cells). Data are means +/- s.e.m. Asterisks denote statistical significance (**p < 0.01, ***p < 0.001) as determined by a two-tailed t-test.

**Supplementary Figure 3.**
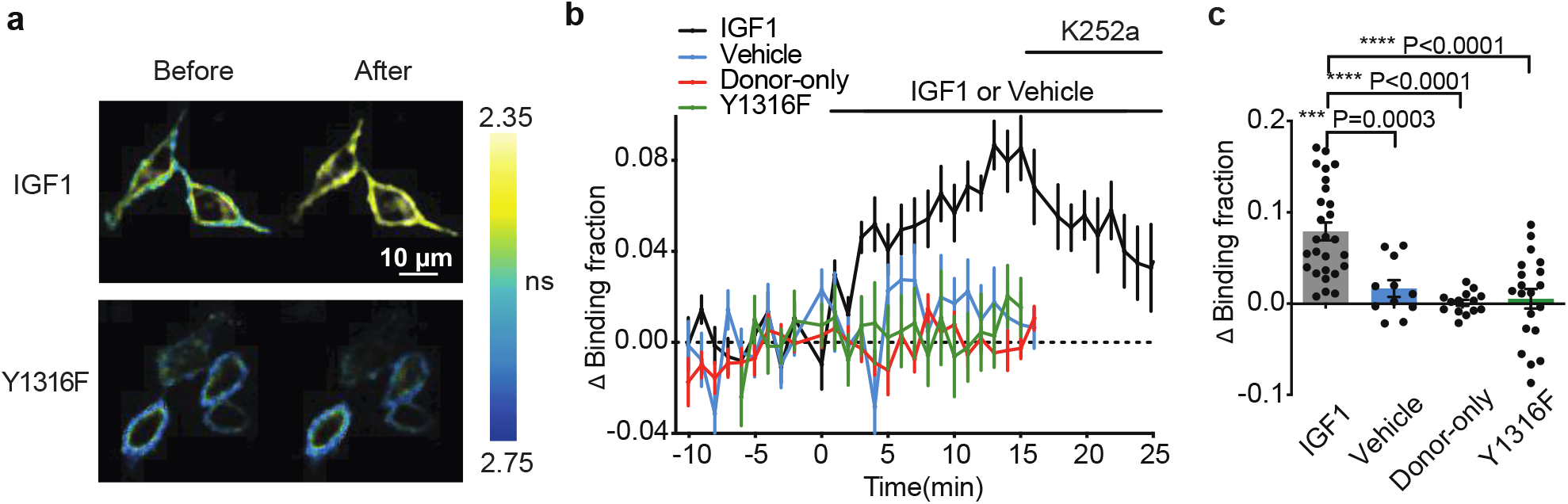
Characterization of IGF1R sensor in HeLa cells. **a**. Representative fluorescence lifetime images of IGF1R sensor expressed in HeLa cells acquired before and 6-14 min after IGF1 (6.5 nM) application (averaged multiple images over 8 min). Shorter lifetime indicates higher IGF1R activity. **b**. Time course of IGF1R activation measured as the change in the binding fraction of IGF1R-mEGFP bound to mCherry-SH2-mCherry before and after IGF1 or vehicle stimulation. For the IGF1 group, we applied a tyrosine kinase inhibitor K252A to the bath at 15 min. In the donor-only group, we expressed only IGF1R-mEGFP. In the Y1316F group, we mutated the IGF1R autophosphorylation site critical for SH2 binding (IGF1R^Y1316F^-mEGFP instead of IGF1R-mEGFP). N = 27/8 for IGF1, 11/4 for Vehicle, 14/5 for donor-only, and 20/6 for Y1316F (cells/experiments). **c**. IGF1R activation (averaged over 6–15min) for experiments in **b**. Data are means +/- s.e.m. Asterisks denote statistical significance (***p < 0.001, ****p < 0.0001) as determined by ANOVA followed by a post-hoc test using Tukey’s correction

**Supplementary Figure 4.**
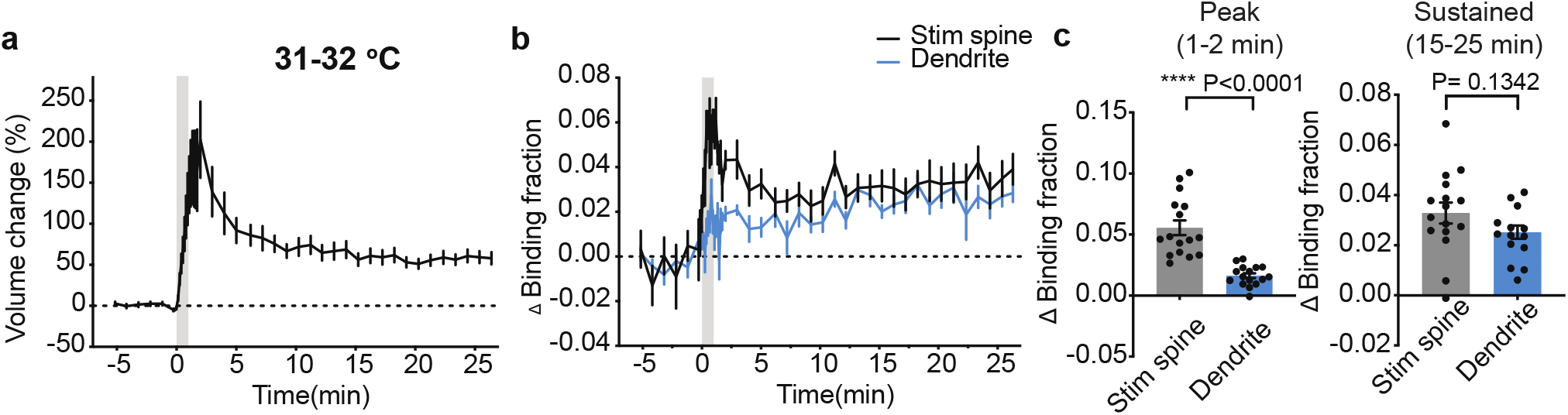
Activation of IGF1R at a near-physiological temperature. Spine volume change (**a**), and time course (**b**) and quantification (**c**) of IGF1R activation in stimulated spines (Stim spine) and their parent dendritic shafts (Dendrite) at 31-32 °C, N = 16/9 (spines/cells). Data are means +/- s.e.m. Asterisks denote statistical significance (****p < 0.0001) as determined by a two-tailed t-test.

**Supplementary Figure 5.**
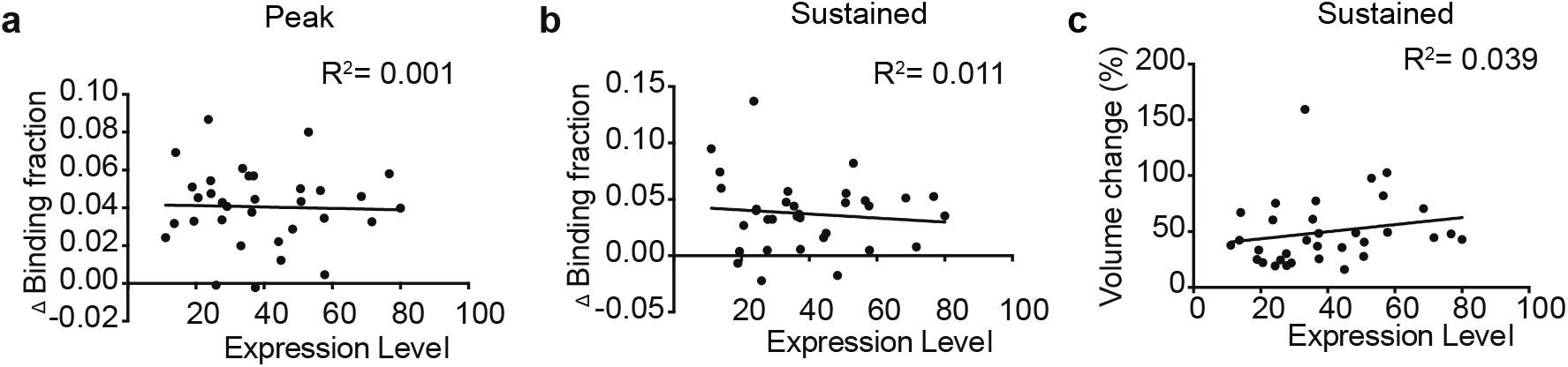
Correlation between IGF1R sensor expression and signal amplitude. IGF1R activations measured as changes in the binding fraction at the peak (1-2 min, **a**) and the sustained phase (15-25 min, **b**) and sustained spine volume changes (**c**, 15-25 min) as a function of relative expression levels of IGF1R donor. We measured the expression level as fluorescence intensity in the primary dendrite. Reanalyzed data from **Figure 2d-h**. The data were fit to a linear regression model with corresponding coefficients of determination (R^2^).

**Supplementary Figure 6.**
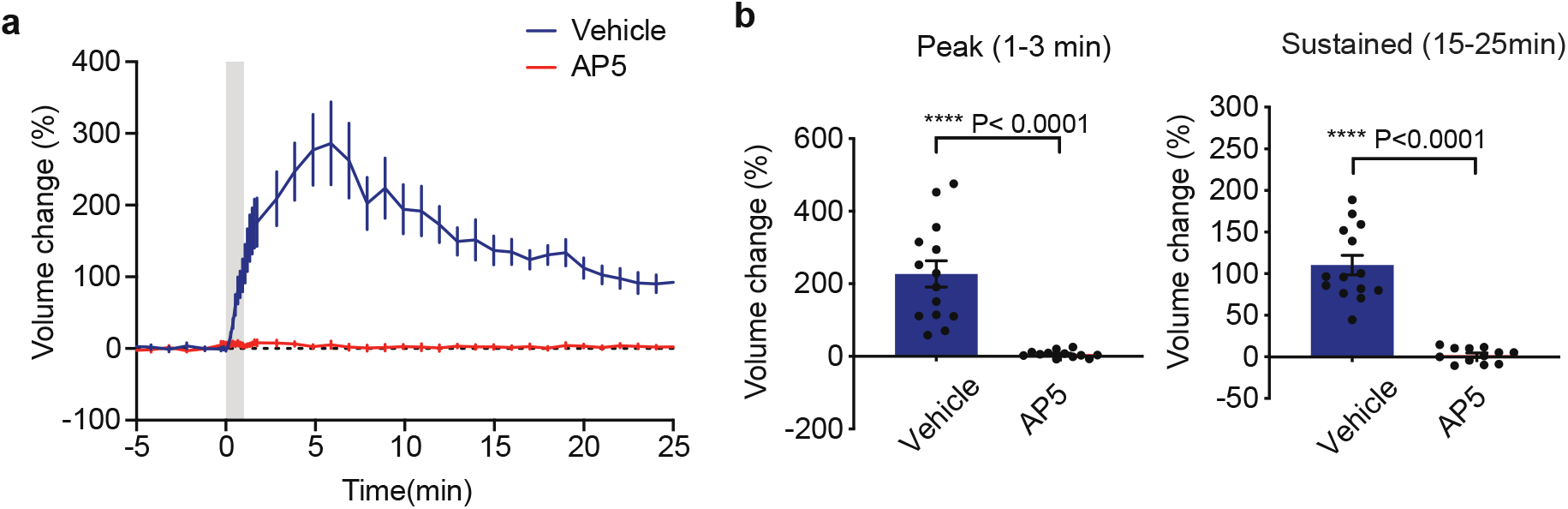
Pharmacology of spine volume changes in neurons expressing IGF1R sensor. Time course (**a**) and quantification (**b**) of spine volume change of spine volume expressing IGF1R sensor in the presence of vehicle (water) or AP5. N = 13/7 vehicle and N= 13/7 for AP5 (spines/cells). Same sample with **Figure 2l, m**. Data are means +/-s.e.m. Asterisks denote statistical significance (****p < 0.001) as determined by a two-tailed t-test.

**Supplementary Figure 7.**
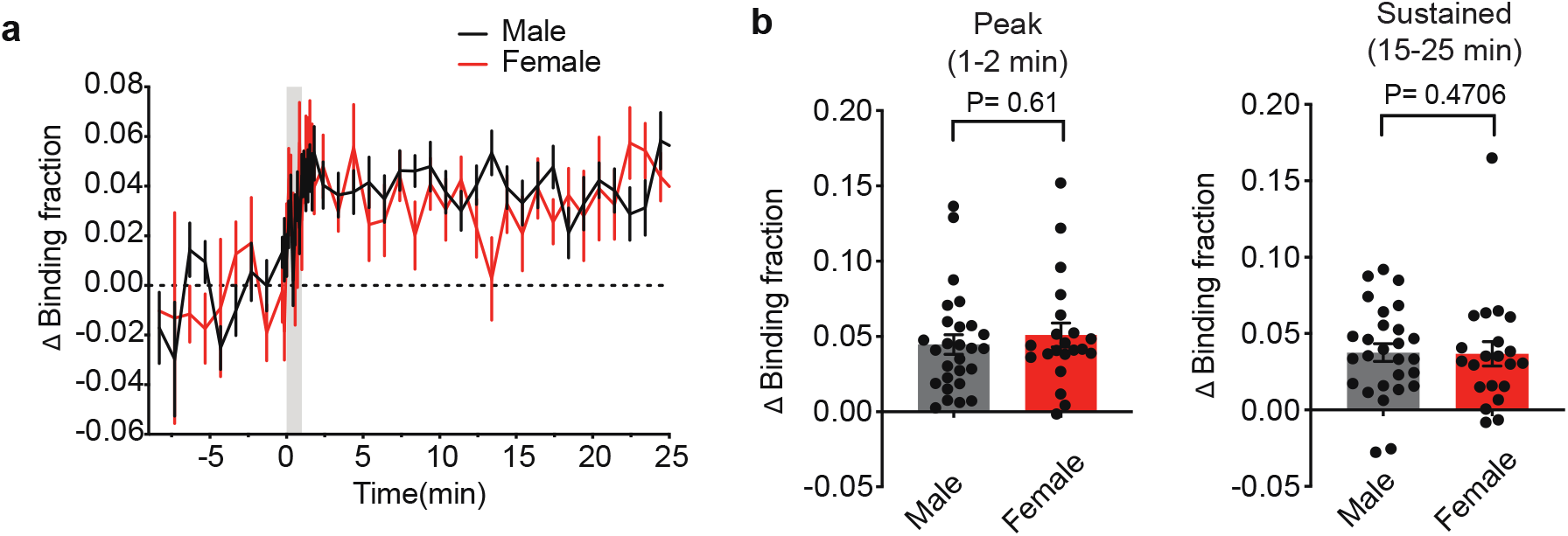
Activation of IGF1R in females and males. Time course (**a**) and quantification (**b**) of IGF1R activation, measured as the binding fraction of the sensor, in stimulated spines and adjacent dendritic shafts in female and male mice. Reanalyzed data from **Figure 2d-h**. N = 27/14 for male and N= 21/11 for female (spines/cells). Data are means +/- s.e.m. P values are determined by a two-tailed t-test.

**Supplementary Figure 8.**
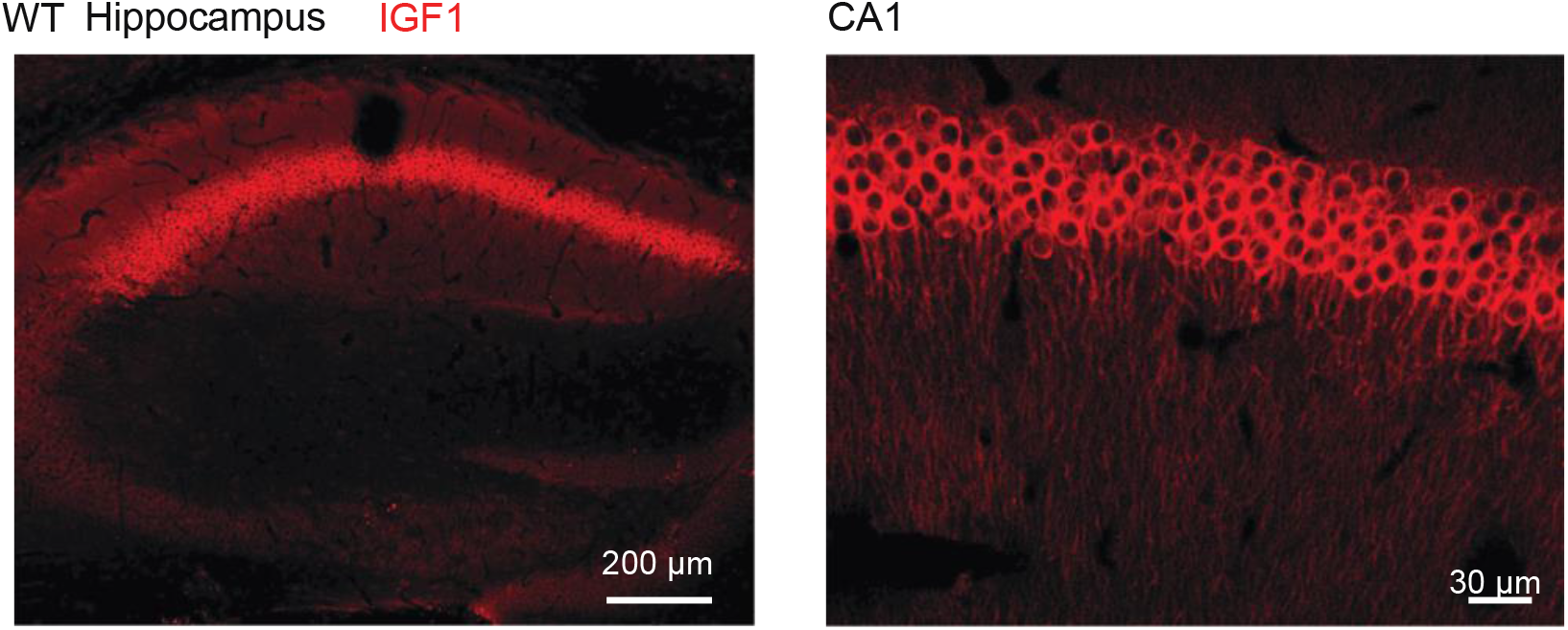
IGF1 expression in the hippocampus of wildtype mice. Representative IGF1 immunofluorescence images of the entire hippocampus (left) and the CA1 region (right) of B6 wildtype mice.

**Supplementary Figure 9.**
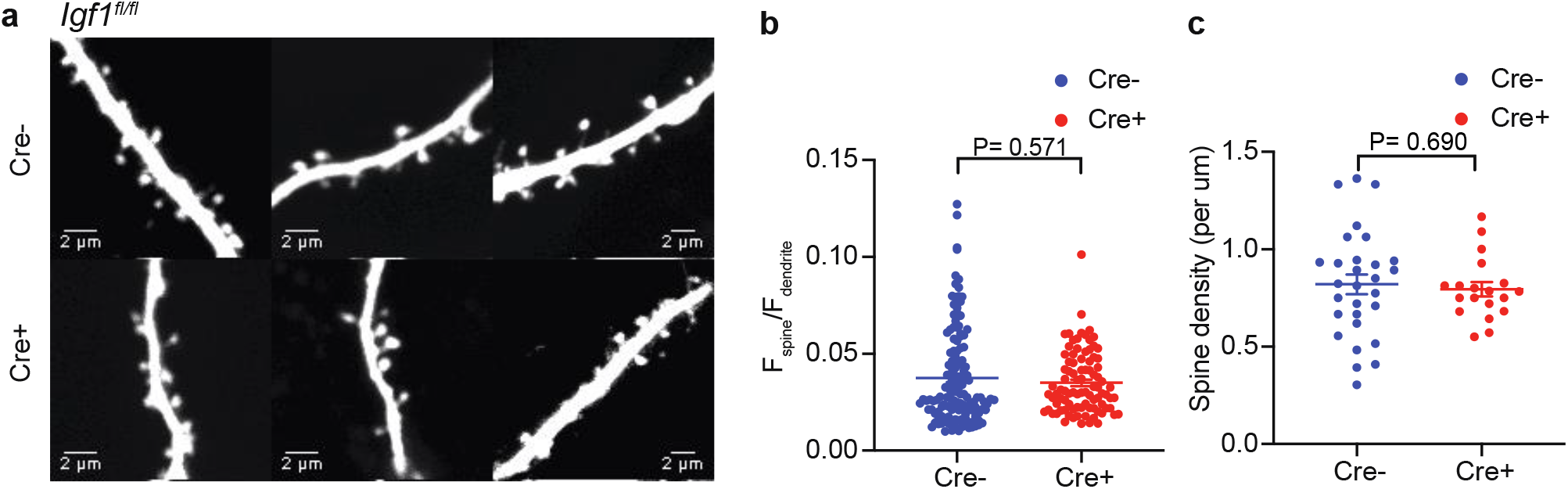
Deletion of IGF1 does not affect basal spine size and density. **a**, Representative 2-photon images of spines in neurons transfected with mEGFP plus tdTomato (Cre-) or mEGFP plus toTomato-Cre (Cre+) for *Igf1*^*fl/fl*^ slices. **b**. The ratio of fluorescence intensity in the spine and that in the dendrite (sum intensity of spine divided by sum intensity of dendrite per 5 µm). N=136/5 for Cre- and N= 91/ 4 for Cre+ (spines/cells). **c**. Density of spines. N=29/5 for Cre- and N= 19/4 for Cre+ (dendrites/cells). Data are means +/- s.e.m. P values are determined by a Mann-Whitney U test (**b**) or a two-tailed t-test (**c**).

**Supplementary Figure 10.**
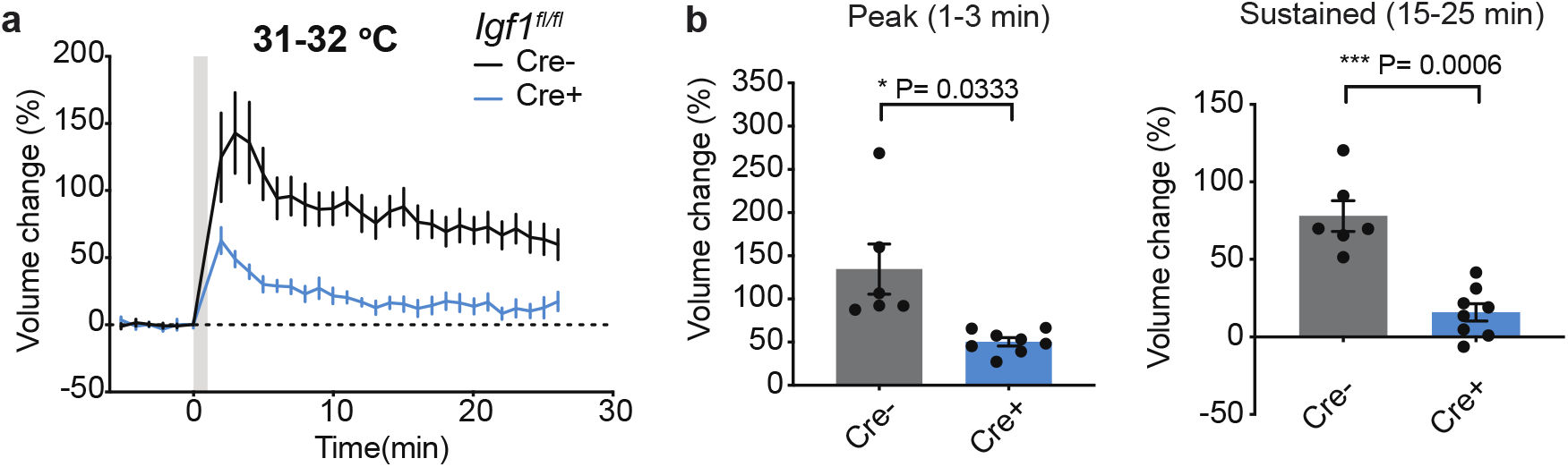
Postsynaptically synthesized IGF1 is required for spine plasticity at a near-physiological temperature. Time course (**a**) and quantification (**b**) of glutamate-uncaging-induced spine volume change for *Igf1*^*fl/fl*^ hippocampal slices in neurons transfected with (Cre+) or without Cre (Cre-) at 31-32 °C. N = 6/4 for Cre- and 8/5 for Cre+. Data are means +/- s.e.m. Asterisks denote statistical significance (*p < 0.05, **p < 0.01) as determined by a two-tailed t-test.

**Supplementary Figure 11.**
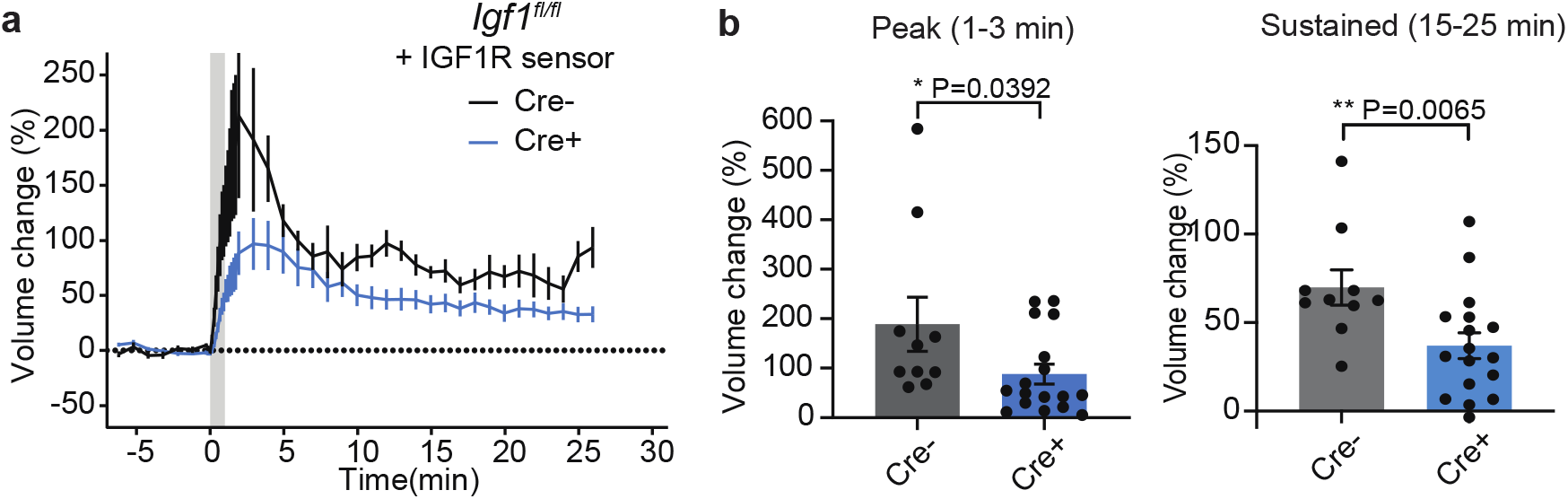
Spine volume changes in neurons expressing IGF1R sensor in IGF1 knockout neurons. Time course (**a**) and quantification (**b**) of glutamate-uncaging-induced spine volume change for *Igf1*^fl/fl^ hippocampal slices in neurons transfected with (Cre+) or without Cre (Cre-). N = 10/6 for Cre- and 16/8 for Cre+. Same sample with **Figure 4c, d**. Data are means +/- s.e.m. Asterisks denote statistical significance (*p < 0.05, **p < 0.01) as determined by a two-tailed t-test.

## Notes

### Competing Interest Statement

RY is a founder of Florida Lifetime Imaging, a company that helps people set up fluorescence lifetime imaging.

